# Vortex light field microscopy: 3D spectral single-molecule imaging with a twist

**DOI:** 10.1101/2024.07.18.604091

**Authors:** Boya Zhang, Sam Daly, Chengxi Zhu, Martin O. Lenz, Lucien E. Weiss, Lisa-Maria Needham, Ruby Peters, Steven F. Lee, Kevin O’Holleran

## Abstract

We introduce vortex light field microscopy (VLFM), a novel method for snapshot 3D spectral single-molecule localization microscopy. Inspired by the azimuthal phase profile of optical vortices, we place an azimuthally oriented prism array immediately after the microlens array in a Fourier light field microscope (FLFM). This innovative arrangement causes the axial position and spectral peak for a point emitter to be encoded in the radial and azimuthal displacement of point-spread-function (PSF) respectively. This enables simultaneous detection of 3D position and emission peak of individual fluorophores with 25 nm spatial precision and 3 nm spectral precision over a 4 *μ*m depth of field (DOF). We illustrate the spectral scalability of our method by performing four-color 3D single particle tracking of freely diffusing fluorescent beads, and two-color 3D dSTORM imaging of microtubules and mitochondria in fixed COS-7 cells, without the need for spectrally distinct fluorophores.

## Introduction

Single-molecule localization microscopy (SMLM) (1–3) is a powerful technique that enables the visualization of various cellular structures at nanoscale. Expanding SMLM with three-dimensional (3D) techniques (4–11) yields deeper insights into the 3D architectures of biological structures. By combining 3D SMLM and multicolor methods, multiple fluorescent labels within the same volume can be imaged simultaneously, which facilitates the study of complex biological phenomena involving interactions between different molecular species.

Various multicolor SMLM methods have been reported in recent decades, such as multiplexing (12–14), spectral demixing (15–17), sequential imaging (18–20) and laser switching (21–23). However, most of these methods are discrete approaches that only allow visualization of a few different molecular species or, for demixing approaches, use molecules with overlapping spectra. Therefore, these are not scalable to the broader spectrum and necessitate prior knowledge of emitters’ spectra. Dispersive and diffractive methods (24–29) offer continuous spectrum measurement with high spectral resolution but spread the PSF over a large number of pixels, which decreases both the signal-to-noise ratio and observable density. Moreover, their 3D information was achieved by incorporating a cylindrical lens to produce an astigmatic 3D PSF (4, 30) into the spatial channel, which has a limited DOF ∼1 *μ*m. Thus, most 3D spectral (*x, y, z, λ*) methods either achieve continuous wavelength with 3D over a narrow axial range (24, 31), or a larger axial range at several discrete wavelengths (14, 32), but not deep 3D position plus continuous wavelength.

To address these limitations, we developed vortex light field microscopy (VLFM), a simple and efficient 3D spectral single-molecule imaging method for the simultaneous detection of 3D spatial position and peak spectral emission of individual fluorescent molecules. The ability to image continuously in a 3D volume is a key benefit from our previous work on single molecule light field microscopy (SMLFM) (10). SMLFM features a PSF composed of an array of Gaussian-like spots whose radial displacement is used to determine the axial emitter location. The primary advantages of SMLFM are large DOF and robustness to aberration, making it particularly suitable for high-density volumetric imaging (33). We have now advanced this method to include a full spectral capability without compromising efficiency or simplicity. This is achieved by encoding spectral information into the azimuthal disparity: a shift in spectral emission peak causes the global PSF to rotate around the optical axis. This is entirely compatible with the existing spatial encoding of fluorescence emitter location which is determined through translation and radial disparity. This compatibility allows VLFM to image over a broad range of wavelengths while maintaining the key advantages of SMLFM. Our method has a simple and compact PSF footprint which is desirable for high-density imaging, and does not require extracting the position and wavelength from a complex PSF model (11). Our method also features optical simplicity, more friendly and cost-effective than methods using multiple paths (14) or dual-objective (24, 31).

In this work, we introduced the principle of vortex light field microscopy along with a guide for its optical design. We quantitatively measured the precision of our method by imaging fluorescent beads, where we achieved ∼20 nm spatial precision and ∼2 nm spectral precision over a DOF of 4 *μ*m. Then we demonstrated our method by simultaneously tracking four spectrally distinct populations of diffusing beads over 4 *μ*m DOF. Finally we applied our method to the imaging of 3D nanoscale architecture of the microtubule and mitochondria network in COS-7 cells, achieving 3D imaging of two dyes 25 nm apart in emission spectrum over a DOF of 4 *μ*m. Overall, our method is robust for quantitative 3D multicolor single-molecule imaging and presents a significant advance for the life sciences.

### Principle of Vortex Light Field Microscopy

A standard widefield microscope can be converted into vortex light field microscopy by placing a microlens array (MLA) and a prism array on the Fourier plane (Fig. 1(a)). These two optical components are the key to achieving 3D spectral single-molecule imaging. They encode the axial position and emission wavelength into the radial and azimuthal displacement of global PSF respectively. Fig. 1(b) shows the effect of emitter’s 3D position and emission wavelength on image formation. The ability of detecting the axial position comes from the MLA. Similar to the Shack-Hartmann wavefront sensor (34), the MLA samples the spatial and angular information of the exit pupil wavefront. A single emitter in the object space will form a PSF composed of an array of spots on the image plane. The center of each focal spot can be localized with a level of precision much smaller than the diffraction limit using current algorithms (35, 36). The disparity for localizations in each perspective view is a function of the average wavefront gradient over the local microlens area. For an axially displaced emitter, it will generate a spherical wavefront whose gradient varies as a function of the axial position, resulting in radial displacements of the PSF. Therefore, the wavefront information are recorded in the position of spot centroids, allowing the PSF to be measured across an extended axial range. As the radial direction has already been used for encoding the emitter’s axial position, we explored the phase profile of an optical vortex to make use of the previously untapped azimuthal direction to encode spectral information. Spectral encoding can be achieved through the diffraction or dispersion of light, with various implementation methods such as spatial light modulators (SLM), diffractive gratings and prism arrays. Among these options, we choose prism array for higher efficiency. The prism array disperses the light chromatically and induces spectral displacement on the PSF. We intentionally choose a small prism angle and low dispersive glass to keep the PSF remaining in compact spots.

**Fig. 1.**
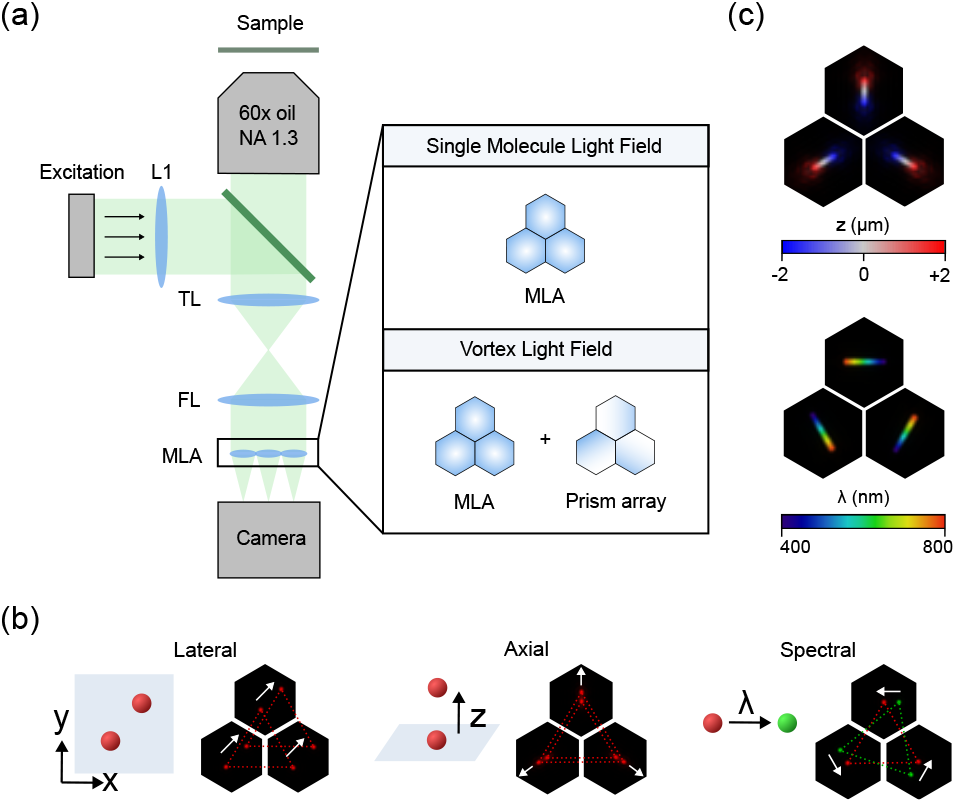
(a) Optical layout of vortex light field microscopy: an extra prism array is added to the Fourier plane, compared to the single molecule light field setup. (b) Simulated PSFs of molecules with different spatial position and emission wave-length. Localizations in each view generated by the same emitter are connected to form an equilateral triangle. Lateral and axial displacements result in the translation and expansion of triangles respectively, while spectral changes result in rotation of the triangle. Note that the triangles are simply a visual aid for comprehension of the principle of VLFM and are not used for reconstruction. (c) Axial and spectral stack projection of simulated PSFs through the customized MLA and prism array.

**Fig. 2.**
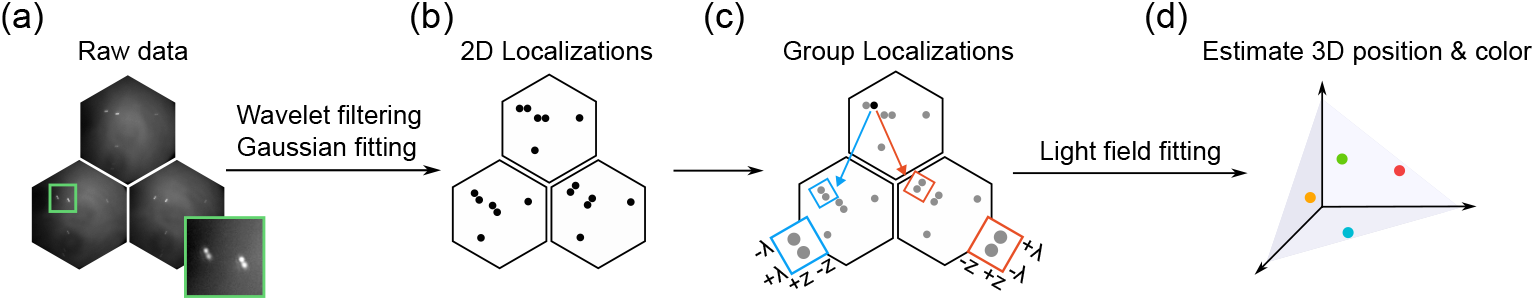
Reconstruction workflow for vortex light field microscopy. (a-b) Point emitters are detected and localized from raw data using wavelet filtering and Gaussian fitting. (c) Localizations in different views that belong to the same emitter are identified and grouped according to z and wave-length constraints. (d) Grouped localizations are fed into the vortex light field model to estimate the 3D position and color of emitters.

## Materials and Methods

### System design

The two key elements involved in the design of VLFM are the MLA and the prism array. Ideally they should be made on the same piece of glass, but due to physical fabrication limits we split them into two parts and place them as close to each other as physically possible. Comprehensive knowledge of the imaging sample is required prior to system design, particularly the sample size and the effective fluorescence emission spectra. The sample size involves the lateral and axial dimensions of the cell, which reflect the required field-of-view (FOV) and DOF respectively. The knowledge of emission spectra refers to the spectra detected by the camera, which is related to the fluorescent probes and other optical components in the system. Together these two factors determine the spectral emission peak and are important to the estimation of required spectral precision.

The spatial performance of VLFM is dictated by properties of the MLA. The design process of a suitable MLA usually starts with the number of microlenses *N*_MLA_ as it is highly related to the DOF. The DOF can be estimated from the full width of the axial PSF (37),

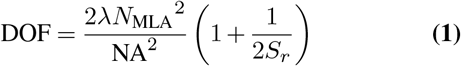

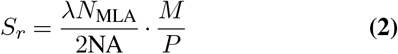

where *λ* is the emission wavelength, NA is the numerical aperture of objective lens, *S*_*r*_ is the pixel sampling rate, *M* is the total magnification and *P* is the camera pixel size. The theoretical calculation matches well with experimental measurements. Since the number of detected photons in each microlens decrease with *N*_MLA_, it is essential to choose the *N*_MLA_ that matches the axial range of the imaging sample optimally. The fill factor, which is the ratio of the back focal plane (BFP) occupied by the MLA, also needs to be considered in the planning of optical configurations. Because the light in the remaining area will generate several cropped views, which cannot be used for localization and result in photon loss. The lens pitch can be easily calculated from BFP size and *N*_MLA_. The total magnification *M* is calculated as the ratio between camera pixel size *P* and Nyquist rate *F*_*s*_, where 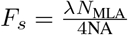 (half of the lateral resolution). The MLA focal length *f*_MLA_ can be derived from the focal length of other lenses and the total magnification *M*.

The prism array determines the spectral performance of our system. Each prism is designed to have the same prism angle but oriented along different azimuthal directions when they are assembled into an array. When we choose the prism material, the refractive index *n*_*d*_ and the Abbe number *V*_*d*_ of the glass need to be considered carefully. Spectral disparity is increased by larger prism angle and using glass with lower Abbe number. However, glass of low Abbe number exhibits larger chromatic aberration and the PSF shape may not stay as compact spot when the prism angle is too large. The refractive index, along with the prism angle, determines the refraction angle and the position of views in image plane. It is essential to keep all the views within the camera’s sensor area.

### Experimental setup

Our VLFM system is based on a standard inverted microscope frame (Nikon Eclipse Ti2) outfitted with a 60×, 1.30 NA infinity corrected, oil immersion objective (Olympus UPlanSApo UPLSAPO60XS2) and tube lens with focal length 200 mm. A 125 mm Fourier lens is placed outside the microscope frame and after the tube lens to form a 4*f* configuration, which relays the back focal plane onto the MLA (f = 68 mm, lens pitch = 1.99 mm, customized from Powerphotonic). The spectral component, hexagonal prism array (N-BK7, pitch = 5 mm, prism angle = 10°, customized from Baiyi Photonics), is held in an XY mount (Thorlabs XYF1/M) tightly followed the MLA. Finally, a Hamamatsu Flash 4.0 V2 sCMOS camera is placed in the focal plane of the MLA for image acquisition. Samples were excited by an Omicron LightHUB (Omicron Laserprodukte Germany) equipped with three lasers (488, 561, 638 nm). Separation between excitation and emission was achieved by a quadband dichroic (Chroma ZT405/488/561/640rpcv2). For cell experiments, the axial focus stabilization was achieved by the integrated Nikon Perfect Focus System.

### Reconstruction

First, a list of 2D localizations is extracted from raw camera images using wavelet filtering and 2D Gaussian fitting. Then the 2D localizations are assigned to each microlens by distance. In the grouping process, localizations correspond to the same fluorescent molecule are identified and grouped for the next calculations. The grouped localizations are fitted using the following equation by least square method to estimate the 3D position (*x*_*i*_, *y*_*i*_, *z*_*i*_) and averaged emission wavelength *λ*,

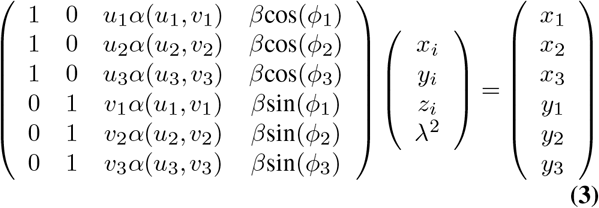

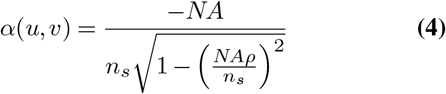

where *α* and *β* are the disparity terms caused by emitter’s axial position and emission wavelength respectively. *ρ* is the radial distance in the normalized pupil coordinates. *uα*(*u, v*) and *vα*(*u, v*) are numerically averaged over the normalized pupil coordinates (*u, v*) for each microlens. *β* is related to prism parameters and obtained by spectral calibration. *φ*_*k*_ is the azimuthal angle of the local prism. (*x*_1_ *x*_2_ *x*_3_, *y*_1_ *y*_2_ *y*_3_) are the relative *x* and *y* distance between 2D localizations and their corresponding microlens centers.

## Results and Discussion

### System Characterization

To quantify the performance of VLFM, we imaged 100 nm fluorescent beads (F8801, Thermofisher) distributed in PBS and immoblized on a #1.5H coverslip. The relationship between precision and number of detected photons was measured on the same set of beads by varying exposure time and applying different neutral density filters to change the range of detected photons. The precision is calculated as the standard deviation of fitted localization over 50 repeats. The precision error is calculated as the standard deviation of precision over multiple beads. In this peformance test using stationary and bright emitters, the axial precision was slightly worse than the lateral precision (∼8 nm). The precision floors of the system are approximately 7 nm and 1 nm for spatial and spectral precision respectively. For detected photons ∼2000, we expect to have 12 nm and 22 nm for lateral and axial precision, and 2.5 nm for spectral precision. Fig. 3(c) and (d) show the precision as a function of the axial position for detected photons up to 6000. The precision was calculated as the standard deviation of fitted values at each 50 nm step with 50 repeats and an exposure time of 10 ms. Spatial precision below 25 nm and spectral precision below 3 nm are witnessed through an axial range of 4 *μ*m.

**Fig. 3.**
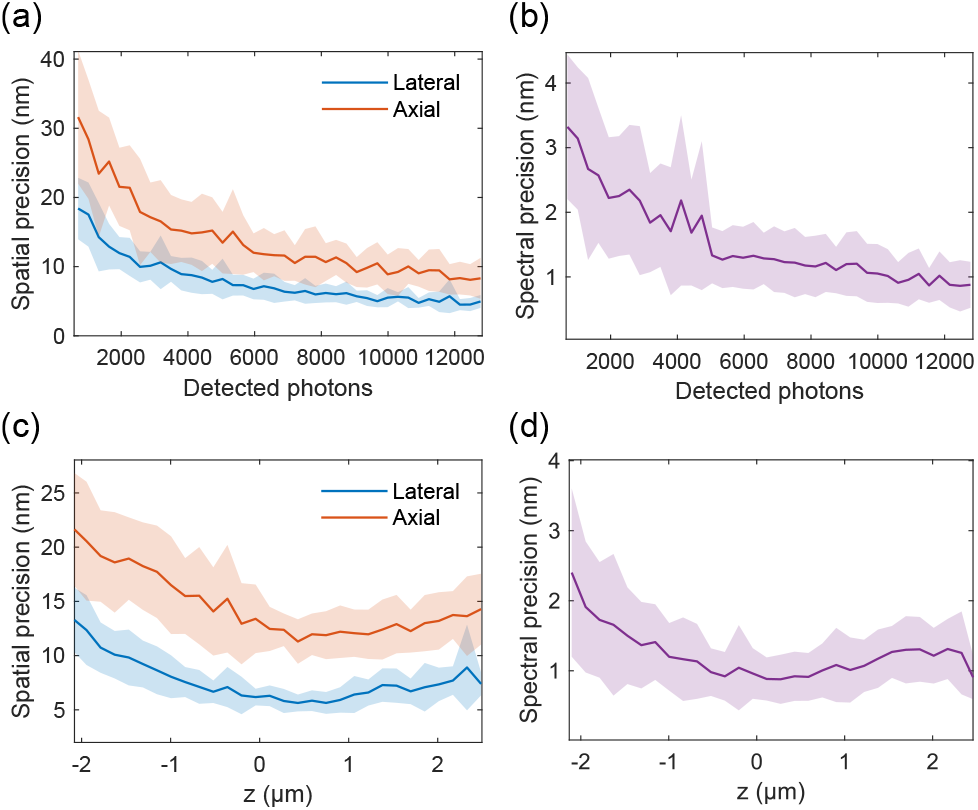
Precision measurements using fluorescent beads (shaded regions represent the precision error). (a) Lateral and axial precision as a function of detected photons. (b) Spectral precision as a function of detected photons. Localizations in (a) and (b) are near the focus plane. (c) Lateral and axial precision as a function of the axial position *z*. (d) Spectral precision as a function of the axial position *z*.

**Fig. 4.**
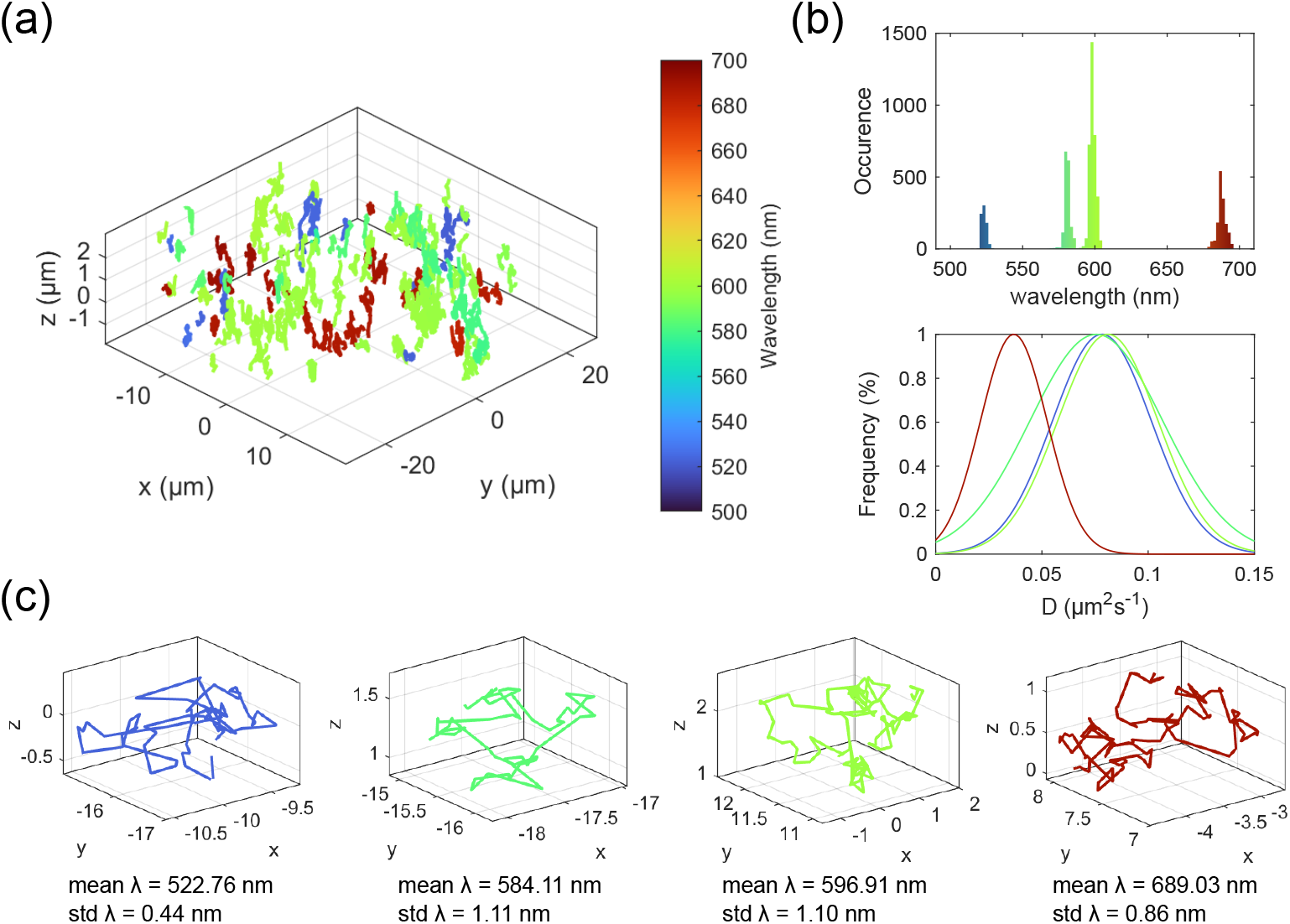
4-color 3D SPT of freely diffusing beads. (a) Rendered 3D trajectories, color-coded by wavelength. (b) Histogram of fitted wavelength for all trajectory points (top) and Gaussian-fitted normalized frequency distribution of measured diffusion coefficient for each bead population (bottom). (c) Examples of isolated trajectory for each bead type.

### Four-color 3D single particle tracking

To demonstrate VLFM’s capability for dynamic 3D + wavelength tracking, we imaged freely diffusing sub-diffraction fluorescent beads with four distinct emission spectra spanning over the visible range. Their emission peaks were at 522 nm, 577 nm, 604 nm and 685 nm respectively. These beads were 100 nm in diameter except for the far-red one which was 200 nm. The beads were diffused in a water-glycerol mixture and simultaneously imaged with 20 ms exposure. 133 trajectories were found from the reconstruction result of 1500 frames. Four distinct peaks were observable in our data, corresponding to wavelengths at 523, 579, 598 and 686 nm with standard deviations of 1.68 nm, 2.02 nm, 2.05 nm, 2.96 nm measured directly from wavelength fits across discrete tracks. Here we showed instant 3D spectral localization with just four fluorescent beads, however with careful selection of dyes and sufficient efficiency our approach could utilize as many as 20 fluorescent probes. This estimate is based on an average peak localization of 10 nm and a spectral range from 400 to 700 nm, including 4 spectral blind spots necessary to block the 4 excitation light sources used in our setup. Of course, using probes with narrow emission spectra and a single excitation source, namely, quantum dots would enable an even larger number of emitters, ∼50, to be monitored simultaneously.

### 3D multicolor dSTORM imaging

Next, we demonstrated the single-molecule sensitivity of our approach by performing direct-stochastic optical reconstruction microscopy (dSTORM) imaging of the microtubule and mictochondrial networks in fixed COS-7 cells. To do so, we labelled these two intracellular structures with two far-red dyes AF647 and CF680 (for microtubules and mitochondria respectively), with heavily overlapping emission spectrum. Since spatial and spectral information is gathered simultaneously in our set-up, imaging was performed with only one line of excitation (638nm) together with a 647nm long-pass filter installed in the detection path to remove scattered excitation laser light. Cells were imaged under highly inclined and laminated optical sheet (HILO) illumination (38) to obtain a high signal-to-noise ratio. Datasets of ∼15,000 frames and 20 ms exposures were acquired over 15 minutes and reconstructed to yield 306,000 3D localizations (≥160,000 after local density filtering to remove potentially overlapping emitters). Importantly, VLFM allows for a significantly larger DOF compared to previous 3D multicolor SMLM methods(16, 24, 31, 39), allowing us to interrogate the structure of biological samples at higher depths while distinguishing the two dyes in the wavelength color-coded image (Fig. S7) without prior knowledge of spectra. To resolve the two structures separately, we categorized each molecule into its respective channels according to a threshold based on the observed wavelength distribution shown in Fig. 5(d). Cross-sections in the xz plane (Fig. 5(c)) reveal the relative 3D positions of the two structures, where mitochondria are sand-wiched between microtubules.

**Fig. 5.**
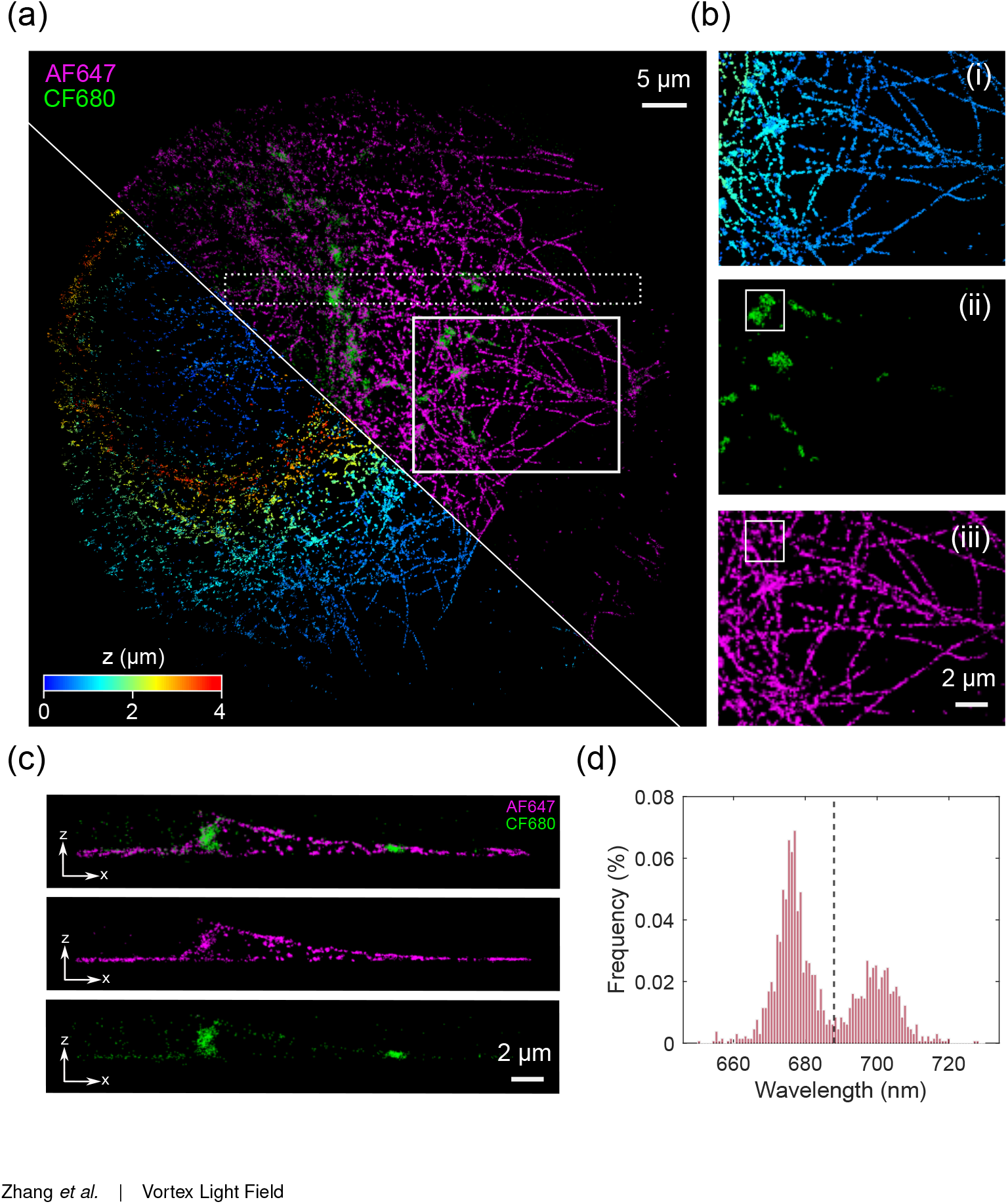
3D two-color dSTORM image by vortex light field microscopy. (a) Composite depth (jet colourmap - left) and spectral (magenta/green - right) color-coded 3D reconstruction of microtubules (AF647) and mitochondria (CF680) in a fixed COS-7 cell. (b) Separated channels for the white region of interest (ROI) in (a); (i) 3D render of all molecules, (ii) molecules assigned to CF680 (thresholded by > 688 nm) and (iii) molecules assigned to AF647 (thresholded by < 688nm). (c) Example vertical xz slice showing microtubules in contact with the top and bottom of mitochondria (dotted region from cell depicted in (a)). (d) Wavelength distribution of the white ROI in (b). Threshold line is set at 688 nm to assign molecules to either the AF647 or CF680 category for visualization.

## Conclusions

Here, we have developed vortex light field microscopy for 3D spectral single-molecule imaging, allowing simultaneous detection of the 3D position and emission peak of individual molecules with a highly efficient system and spatially compact PSF. Importantly, in this technique, spatial and spectral information (*x, y, z, λ*) can be extracted from 2D localizations as our method encodes the axial position and emission wavelength in the radial and axial displacement of PSF respectively. Using fluorescent beads at conditions that approximate single-molecule-level signals and background, the precision was 25 nm spatially and 3 nm spectrally over a DOF of 4 *μ*m for just a few thousand photons. Notably, under highly optimized experimental conditions, some fluorophores can be much brighter (Dempsey), thus this approach represents suitability for a wide range of emitters. We demonstrated dynamic imaging by tracking of four types of freely diffusing beads, showing our technique is applicable to a large spectral bandwidth while achieving excellent spectral precision such that many different emitter types could be deployed simultaneously. This is further evidenced by our multicolor dSTORM experiment, where the emission spectra of our emitters overlapped significantly, yet could still be resolved over a 4 *μ*m axial range.

We estimate that the combination of high spectral precision and dynamic range could potentially handle a large number of fluorescent probes (∼50) to be identified and localized in 3D, which could possibly be coupled with techniques like DNA-PAINT for multiplexed super-resolution, or single-molecule fluorescence in situ hybridization (smFISH) (40) to offer high-throughput RNA imaging, or with nanobarcoding to study spatial proteomics in neurons (41, 42). Our method could also be applied to distinguish multiple spectrally distinct probes that share the same spatial position, such as in single-molecule Förster resonance energy transfer (smFRET) (43, 44), or more simply, colocalization analysis without FRET (45), as it doesn’t necessitate close spatial proximity between the probes. Another potential application is monitoring the spectral fluctuations or environmental sensitivity of fluorescent probes (25), given our method can measure the spectral information continuously.

While a single-molecule imaging implementation makes the analysis straightforward, the sparsity condition of singlemolecule imaging is also particularly well suited to such spatial and spectral information coding; however, in principle, the general concept presented is not just limited to localization microscopy. Our method could be used for simultaneous multicolor microscopy to generate diffraction-limited video frame rate spectral 3D data sets. Future developments could focus on computational methods to further increase localization density, such as better grouping strategies and multi-emitter fitting. Probe designs that allow numerous narrow emission profiles to be simultaneously imaged without buffer changing would also improve the spectral capacity of our method. Robustness to aberrations could be enhanced by integrating the MLA and prism array into one piece of glass. Metasurface and flat optics also offer interesting avenues of exploration to further enhance spectral and spatial sensitivity. Overall, the 3D spectral information coupled with the simple PSF footprint enables vortex light field microscopy to be a promising method for high-precision spectral volumetric imaging.

## ACKNOWLEDGEMENTS

The authors would like to thank Ana Fernandez-Villegas for providing COS-7 cells. B.Z and K.O.H conceptualized the project. K.O.H, R.P and S.F.L supervised the project. K.O.H and L.-M.N administered the project. B.Z and K.O.H developed the methodology. B.Z and K.O.H designed the optics. B.Z, K.O.H, M.O.L and L.E.W built the optical system. B.Z conducted the experimental investigation. B.Z wrote the software and performed simulation and data analysis. R.P designed the cell experiment and optimized the labelling method. R.P and B.Z prepared the dSTORM samples and performed cell experiments. S.D prepared characterization samples and provided a tracking code. C.Z optimized Zemax simulations and tracking experiment. B.Z and K.O.H wrote the paper. All authors reviewed and edited the final paper.

